# A perspective on information optimality in a neural circuit

**DOI:** 10.1101/2021.10.27.466179

**Authors:** Friedman Robert

## Abstract

The nematode worm *Caenorhabditis elegans* has a relatively simple neural system for analysis of information transmission from sensory organ to muscle fiber. Therefore, an example of a neural circuit is analyzed that originates in the nematode worm, and a method is applied for measuring its information flow efficiency by use of a model of logic gates. This model-based approach is useful where the assumptions of a logic gate design are applicable. It is also an useful approach where there are competing mathematical models for explaining the role of a neural circuit since the logic gate model can estimate the computational complexity of a network, and distinguish which of the mathematical models require fewer computations. In addition, for generalization of the concept of information optimality in biological systems, there is an extensive discussion of its role in the genetic-based pathways of organisms.

## 1. Introduction

### 1.1. The logic gate model

McCulloch and Pitts introduced the concept of a logic gate model to explain the processing of information in an animal neuronal system, based on the concept that a neuron is in a resting or active state, and the synaptic connections exhibit specific non-dynamic behaviors [1]. The logic gate design is an efficient model of boolean algebraic operations, including the basic operators AND, OR, and NOT (Fig. 1). However, others consider the biological design of a neuronal network inconsistent with the assumptions of this model [2–12]. Even though the logic gate model is not expected to predict behavior of a neuron at the cellular level, it does offer a measurement of expected computational efficiency in a neural network. Given knowledge of a neural system’s layer of nodes of inputs, a layer of nodes of outputs, the assumption that a neuron is either resting or active, then it is possible to estimate the optimal number of logic gates to model the information optimality and potentially the behavior of the system. This metric of optimality can also be used in comparisons among neural circuits, and potentially as a source of intuition on their network structure and function.

**Figure 1.**
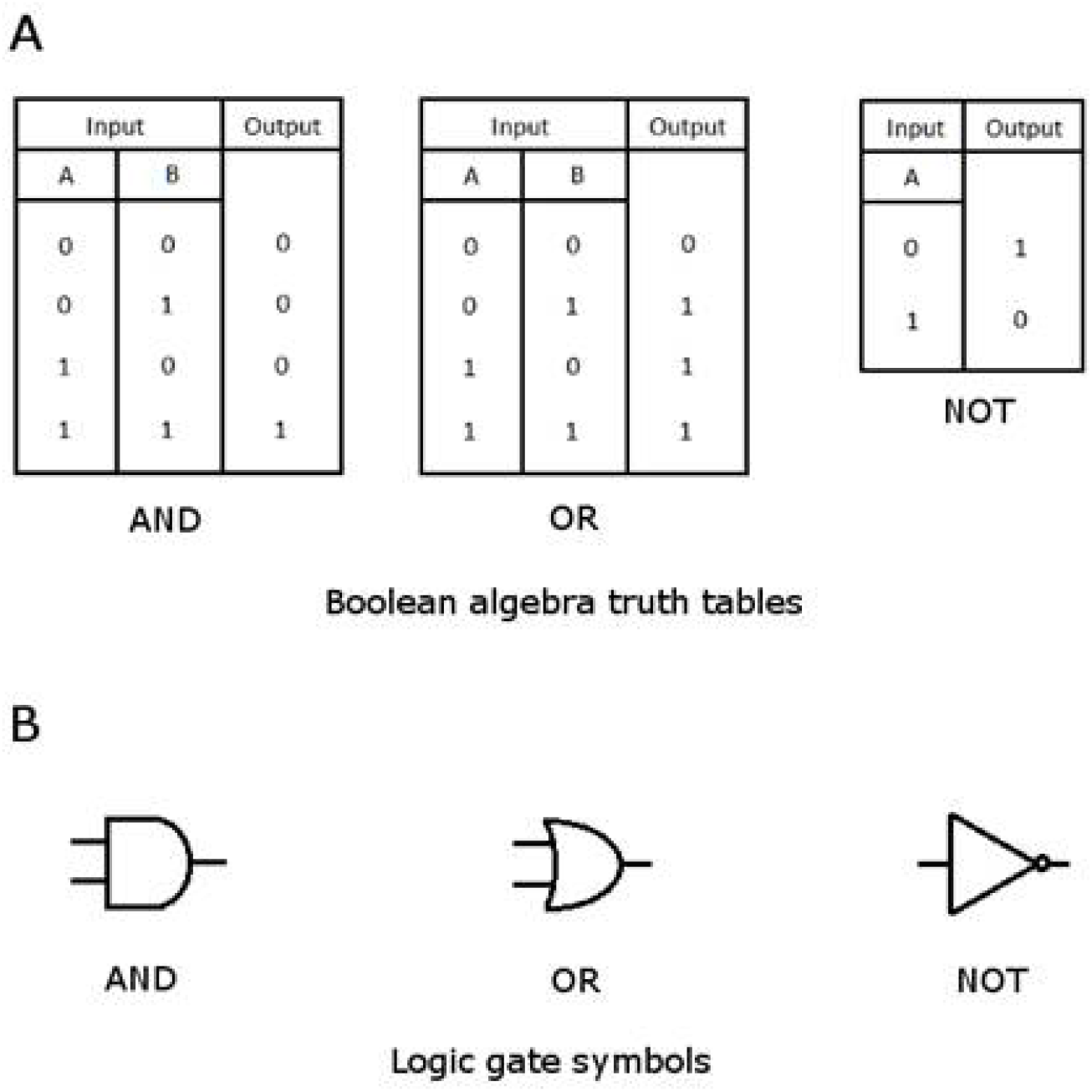
A. Truth table for the basic boolean operations AND, OR, and NOT. B. The logic gate symbols represent the three basic boolean operations.

The assumption where a neuron as active or at rest can be represented by an assignment to binary values, 1 and 0. Boolean algebra defines logical operations with these binary values, and originally named after the 19th century British mathematician George Boole [13], although the German polymath Gottfried Wilhelm von Leibniz had developed an equivalent mathematical system in 1686 [14–16]. Leibniz had also developed a mechanical system for computing with binary values, an analog of the modern digital computer [16–17].

One problem of modeling neuronal communication is the availability of reliable data on a neural system and its activity. This activity is dependent on external effects, such as the external environment, the interactions between the organism and environment, and spatial and temporal nonlinear dynamics. The complexity of this system is expected to grow exponentially with variation in these environmental conditions, and likewise with organism complexity, including all possible physiological states. This effect is observed in the sociality of ants [18]. It is not likely that a neural system has separate circuits for each environmental condition, so it is expected that a neural circuit is adapted for multiple functions. Modularity is another strategy for enhancing the efficiency in information processing, such as observed in honeybee sociality [19].

Information compression, such as described above where a neural circuit has multiple functions, also occurs in other information-based systems, such as in transmitting audio over a network cable. This property is expected in a network designed for efficiency. This is also a confounding factor for disentangling the set of functions of a neural circuit since it has greater than one function. However, in the case of a simple neural circuit, along with a controlled environment, then it should be possible to gain insight on information processing by mapping a neural circuit to a simple model, and therefore resist the expected effects of the complex dynamics in a neural system.

### 1.2. Efficiency of mathematical operations

It is also possible to consider the efficiency of multiplication and division operations in the context of boolean algebra and logic gates. Hypotheses and models of fly motion detection involve the use of these operations.

Borst and Helmstaedter [20] had reviewed prior work on visual motion detection in the fly. In their review of potential models for motion detection, they discussed a model’s dependence on sampling luminance values at two or more points in a visual space. One mathematical model includes a calculation of velocity, along with a division operation. Another model relies on a computation with a correlation between two spatially separated luminance values. They further discuss that the correlation-based model is the preferred one because it fits better with current knowledge about the information processing in the fly visual system.

Theory and empirical studies led to a later study, and in support of the latter model, otherwise referred to as a “delay-and-compare model” of fly motion detection, where Groschner and others [21] showed evidence of a “multiplication-like nonlinearity” in a single neuron of this system, a prediction expected by prior theory on information flow [21–22]. This process is also shown to be dependent on a biological mechanism involving “pairs of shunting inhibitory and excitatory synapses” [21–22]. This merging of theory with biological evidence has the potential to produce biologically realistic models of neural circuits, with an expectation of applicability across animals, given the level of evolutionary convergence in neural circuitry and similarity in their molecular mechanisms for neuronal communication [20–21].

Even though biologically realistic models are generally preferred, it is still possible to idealize the information flow for making predictions on the optimality of a neural circuit or system, as suggested by this study. If theory predicts a high complexity in the computation of information, regardless of an artificial or biological system, then physical limitation will constrain the nature and performance of the computations. Therefore, this study includes examples from boolean models and their efficiency, including in the case of multiplication and division, operations not native to boolean algebra.

### 1.3. Optimality of a neuronal system

This question of optimality in a neural system has been examined at the level of structure and function in a neural network [8, 23–27]. In the case of the nematode worm *Caenorhabditis elegans*, one type of optimization is observed in the neuronal wiring length for minimizing the path length for information communication [24]. Given a neuronal circuit, it is expected that the system is optimized for the overall cost across development of the animal and the continuing function of the system. In support of this hypothesis is the role of adaptation in the optimization of the anatomy and physiology in animals. Another argument is the role of natural selection that tends to disfavor individuals with inefficient use of its energy, such as evolution of a large central neural system, along with a high energy requirement, in the case where it offers no benefit over the ancestral form. This is wasted energy and results in individuals weakening their chance in the “struggle for existence” against conspecifics.

Other structural properties of the *C. elegans* neural network have been described, such as an observation of a small-world architecture [5, 28–29], sparsity of neuronal connectiveness [11], and commonly observed motifs [3, 30].

The simpler neural systems, such as isolation of a neural circuit, are tractable for measuring information flow efficiency, preferably across an animal’s life history. Past studies have examined the coding of animal behavior in the neural circuits of the nervous system [4, 7, 9, 24, 31–34]. It is further possible to relate these neural systems to information flow in animals [5, 35]. The concept of information flow, efficiency, and the presence of neural circuits suggest an applicability of information flow theory, and use of a logic gate model to estimate the rate of information flow in a system [3, 36]. Other examples of application of artificial models to explain general neural system function includes studies by Yan and others [26], Karbowski [11], and Lysiak and Paszkiel [37]. In particular, Yan and others [26] proposed a model based on control theory to explain and predict the structure of a neural circuit involved in *C. elegans* locomotion. Their predictions were further supported by empirical verification. The prospect of building models at the small and large scale of life is also present in the study of ecological systems, another area involving nonlinear dynamic processes, where the efficiency and flow of energy and matter are measured without parameterizing the properties of the living organisms themselves [38].

## 2. Methods

### 2.1. Data retrieval

The web-based interface at WormAtlas.Org documents the sensory and motor neurons in the adult hermaphroditic form of *C. elegans* [39–44]. This species has two distinct nervous systems, a somatic system of 282 neurons [39] and a pharyngeal system of 20 neurons [40]. Several of these neurons are not fully characterized, and others do not have distinctive membership to either the pharyngeal or somatic system, so it is possible to make assignments based on the curated databases at WormAtlas.Org.

The neural circuit for gentle touch in locomotion is described by Yan and others [26] and Driscoll and Kaplan [31]. This study also assumes that any neural circuit operates by a forward feedback loop, a biologically plausible assertion [30].

### 2.2. Data processing and visualization

Minitab statistical software [45], version 14, provided for data organization and processing. The diagram of the digital circuit was drawn in TinyCAD [46]. It includes software libraries *74TTL* and *gen_Logic* which include graphical symbols for commonly used logic gates.

### 2.2. Logic gate analysis

The Espresso software provides solutions on the minimal number of logic gates given a number of inputs and outputs in a circuit [47]. Below is an example of the input data with 3 inputs and 4 outputs, corresponding to the AND/NOT case (Table 1):

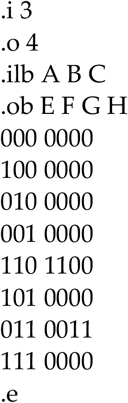

**Table 1.** Truth table for a digital circuit with 3 input and 4 output operations. Each row corresponds to 1 of the 8 possible input and resulting output states. The binary digit 1 is an ON state and 0 is an OFF state. An OR operation resolves to an ON state if either of two specific inputs are ON, while an AND operation resolves to an ON state where two specific inputs are ON. NOT modifies the state from ON to OFF or OFF to ON.

| OR / NOT |  | AND / NOT |  |
| --- | --- | --- | --- |
| Inputs | Outputs | Inputs | Outputs |
| 0 0 0 | 0 0 0 0 | 0 0 0 | 0 0 0 0 |
| 1 0 0 | 1 1 0 0 | 1 0 0 | 0 0 0 0 |
| 0 1 0 | 0 0 0 0 | 0 1 0 | 0 0 0 0 |
| 0 0 1 | 0 0 1 1 | 0 0 1 | 0 0 0 0 |
| 1 1 0 | 1 1 0 0 | 1 1 0 | 1 1 0 0 |
| 1 0 1 | 0 0 0 0 | 1 0 1 | 0 0 0 0 |
| 0 1 1 | 0 0 1 1 | 0 1 1 | 0 0 1 1 |
| 1 1 1 | 0 0 0 0 | 1 1 1 | 0 0 0 0 |

The above list is a computer-readable list of truth table values where each of the lines of input (A, B, C) and output (E, F, G, H) are separated by a space character. The header information includes the count of columns of input and output values, along with a human-readable name for each of the columns of values. The software resolves the truth table into a set of logic gates as represented in boolean algebraic form:

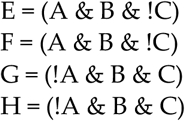

> The software performs the above calculation and solution at a command line: espresso.exe -oeqntott -Dso -S1 -t input_datafile

As the columns of input values increase, the time complexity increases exponentially. For an example of enumerating the input state values in a truth table, a line of bash shell code is below:

> for i in $(seq 0 4095); do echo “obase=2;$i” | bc | tr -d ‘\r’ | xargs printf “%012d\n”; done

## 3. Results

### 3.1. Optimality of an idealized neural circuit

Yan and others [26] examined a neural circuit in *C. elegans* and the sensory neurons involved in gentle touch, the interneurons for control, and the motor neurons that synapse with muscle by neuromuscular junctions [31]. They isolated the essential members of the circuit as predicted by control theory in engineering. The inputs include three sensory neurons, while the outputs correspond to four motor neurons that synapse on locomotory muscles involved in forward and backward motion. Interneurons may act as intermediaries to information flow between sensory and motor neurons.

In this study, the logic gate model was designed for the above number of sensory inputs and motor outputs. Table 1 shows this data as represented by a truth table, and suggests two logic gate models that are abstract representatives of the gentle touch circuit in *C. elegans* [26]. Each 1 or 0 in the truth table corresponds to an ON or OFF state in the neural circuit. The OR/NOT logic gate model allows for multiple neurons to independently code for forward or backward locomotion, while the AND/NOT model requires a neuron to depend on another neuron for any activation of locomotory action. This method is also dependent on the assumption of unidirectional information flow from sensory input to motor output.

Fig. 2 shows a logic gate diagram that corresponds to the solution for the AND/NOT case (Table 1). The diagram also shows the flow of information form sensory input to motor output, along with the connections. As described as priors in the input data (Section 2.2), the number of sensory inputs is 3 and the number of motor outputs is 4. The diagram and model represents an abstract perspective of the neural circuit. The associated procedure can be applied to other neural circuits, too.

**Figure 2.**
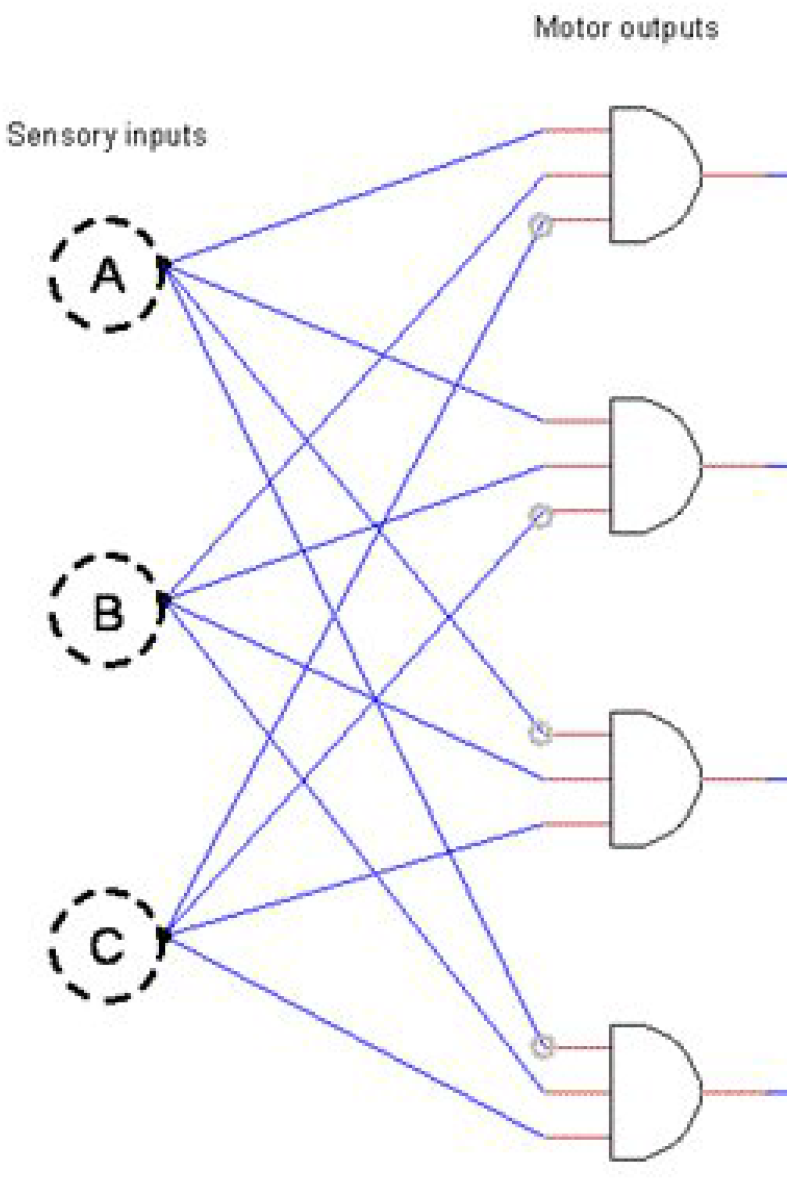
Diagram of a neural circuit with three sensory inputs and four logic gates that correspond to the four motor outputs. The diagram represents a solution for the AND/NOT case in Table 1. The small grey circles between red and blue lines represent a potential inverting of the input signal, a NOT operation.

### 3.2. Efficiency of mathematical operations in a neural circuit

The information-based cost of the operations of multiplication and division provide insight to the processing in an idealized neural circuit. Table 2 lists a case where a 2-bit value is multiplied by another 2-bit value. This is repeated for the case of division to provide a contrast for the cost of these binary operations.

**Table 2.**
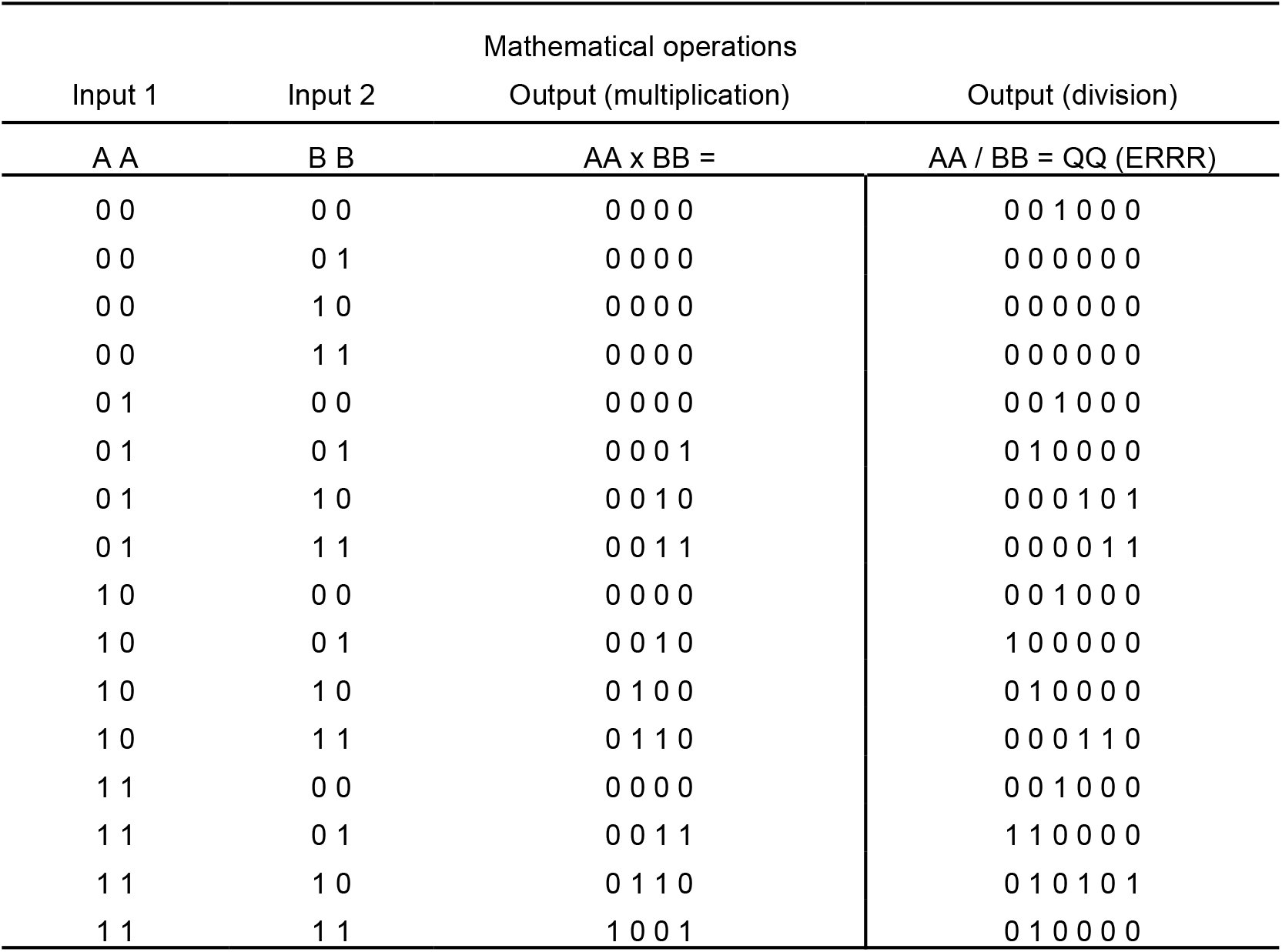
Truth table for binary multiplication and division in the case of two 2-bit input values. Input 1 is a 2-bit value labeled AA; Input 2 is a 2-bit value labeled BB. The output value for multiplication is the product of AA multiplied by BB. The output value for division is AA divided by BB; and, the quotient value QQ is stored in the 2 most significant bits, and the divide-by-zero error bit is the next adjacent bit, while the 3 least significant bits store the unrounded remainder value (RRR). Specific to this case of low precision, the remainder value is a 3-bit value since the highest possible remainder value is 6 in the base 10 number system.

| Input 1 | Input 2 | Mathematical operations |  |
| --- | --- | --- | --- |
|  |  | Output (multiplication) | Output (division) |
| A A | B B | AA x BB = | AA / BB = QQ (ERRR) |
| 0 0 | 0 0 | 0 0 0 0 | 0 0 1 0 0 0 |
| 0 0 | 0 1 | 0 0 0 0 | 0 0 0 0 0 0 |
| 0 0 | 1 0 | 0 0 0 0 | 0 0 0 0 0 0 |
| 0 0 | 1 1 | 0 0 0 0 | 0 0 0 0 0 0 |
| 0 1 | 0 0 | 0 0 0 0 | 0 0 1 0 0 0 |
| 0 1 | 0 1 | 0 0 0 1 | 0 1 0 0 0 0 |
| 0 1 | 1 0 | 0 0 1 0 | 0 0 0 1 0 1 |
| 0 1 | 1 1 | 0 0 1 1 | 0 0 0 0 1 1 |
| 1 0 | 0 0 | 0 0 0 0 | 0 0 1 0 0 0 |
| 1 0 | 0 1 | 0 0 1 0 | 1 0 0 0 0 0 |
| 1 0 | 1 0 | 0 1 0 0 | 0 1 0 0 0 0 |
| 1 0 | 1 1 | 0 1 1 0 | 0 0 0 1 1 0 |
| 1 1 | 0 0 | 0 0 0 0 | 0 0 1 0 0 0 |
| 1 1 | 0 1 | 0 0 1 1 | 1 1 0 0 0 0 |
| 1 1 | 1 0 | 0 1 1 0 | 0 1 0 1 0 1 |
| 1 1 | 1 1 | 1 0 0 1 | 0 1 0 0 0 0 |

The truth table for binary multiplication and division (Table 2) was formatted as an input file for the Espresso software [47]. The multiplication operation has a cost of 8 boolean operations, while division has a cost of 12. This result is dependent on the precision in the remainder value, so increasing this value leads to a higher number of operations. This shows a procedure for use in comparing prior mathematical hypotheses that reflect the kind of computation that is expected in a neural circuit. It also offers a method for estimating the general complexity of information processing in a neural circuit. In reference to computational complexity, the above result supports that multiplication is expected to be favored over division in a system based on information flow, and for a biological case where these neural circuit operations are applicable and plausible.

## 4. Discussion

### 4.1. The utility of a logic gate model

Fig. 2 shows a logic gate model for an idealized neural circuit. This assumes a level of abstraction of the individual parts of the system, and a static snapshot of neuronal dynamics. In a large network, such as for advanced information processing, it is expected that the use of nonlinear dynamics will better capture the deeper complexity of the system, such as used in comparisons of an animal’s neural network to an artificial analog [48–49]. However, at these scales, there is typically a limitation on explainability by a model, particularly at the neuronal level. Instead, use of a logic gate model may provide explainability and intuition on the routing and efficiency of information in a neural network.

The logic gate model is also more applicable for a neural circuit in an unchanging environment, both internal and exernal to an animal. For a dynamic environment, then it becomes difficult to disentangle the inner workings of a neural circuit because it is expected that many of its functions are unknown. It is a reminder that past studies have identified roles of neurons in a controlled setting, however, these roles are commonly specific to one environment, or a few environments, and not tested across all possible environments. This problem is compounded by the complexity inherent in the dynamics of an animal’s physiology and its interactions. This complexity may also be described as a system with high dimensionality, so that the number of parameters is large and difficult to estimate in practice.

The use of the logic gate model depends on information flowing from sensory input to output, such as in the case of a sensory neuron to a motor neuron [50]. The organization of a neural network and its circuits are expected to depend on the values at the input and output layers, especially during the rapid modifications over an animal’s early development. These systems are essentially a programmable network system of neurons and synaptic connections. The connections, at least as idealized features, are dynamic with respect to gain, loss, and in the strength of connection. The connectivity and the network structure show evidence of optimality in structure, as described in the Introduction, and so it is reasonable to consider optimality in the information flow across a neural system, too. Theory provides an important guide, while the above method offers an alternative method where theory is not applicable or easily interpretable.

### 4.2. Information processing as an algorithm in biological systems

#### 4.2.1. Overview of information-based systems

The logic gate model applies to the information processing of logical operations. While it is based on a binary number system, it provides general insight into information flow and routing. The reason is that information-based systems in Nature, in particular the evolved systems of biological organisms, are typically an optimal solution within the constraints of their biological design. Further, the processing of information in these systems is in essence an algorithm and a model. This description of information processing is also present in the process of genetic inheritance, neural systems in biology, and detection of intracellular pathogens in jawed vertebrates.

An animal neural system processes information, and the operations of information processing is in essence an algorithm. The algorithm is stored in the neural network, so it is expected to show a complex data structure, and is contained in a higher dimensional space than expected from the commonly used meaning of *algorithm*. This system, along with other examples in Nature, are described below.

#### 4.2.2. Genetic inheritance of protein sequences

It is possible to frame the inheritance of genetic material using the above perspective (Fig. 3). The genetic material is a sequence of elements, where each element is a nucleic acid molecule, and is inherited by an individual, the progeny, for coding the information and to for a new organism. These processes are usually presented with terminology and pathways that originate from the chaotic path of scientific discovery, and a pre-Linnaean approach to ontology of natural systems, but these informational processes are generalizable and categorical. In the case of generating proteins, the above genetic sequence is the informational template that is processed for creating multiple copies of itself, and then processed further to convert the copies to molecular sequences of amino acids, a protein sequence. These proteins then fold to form complex shapes in three dimensional space. This higher dimensional form leads to complex dynamics, such as its interactions with other molecules, its potential to increase the rate of biochemical reactions, and transforming of stored energy in molecular bonds to physical motion at the molecular level.

**Figure 3.**
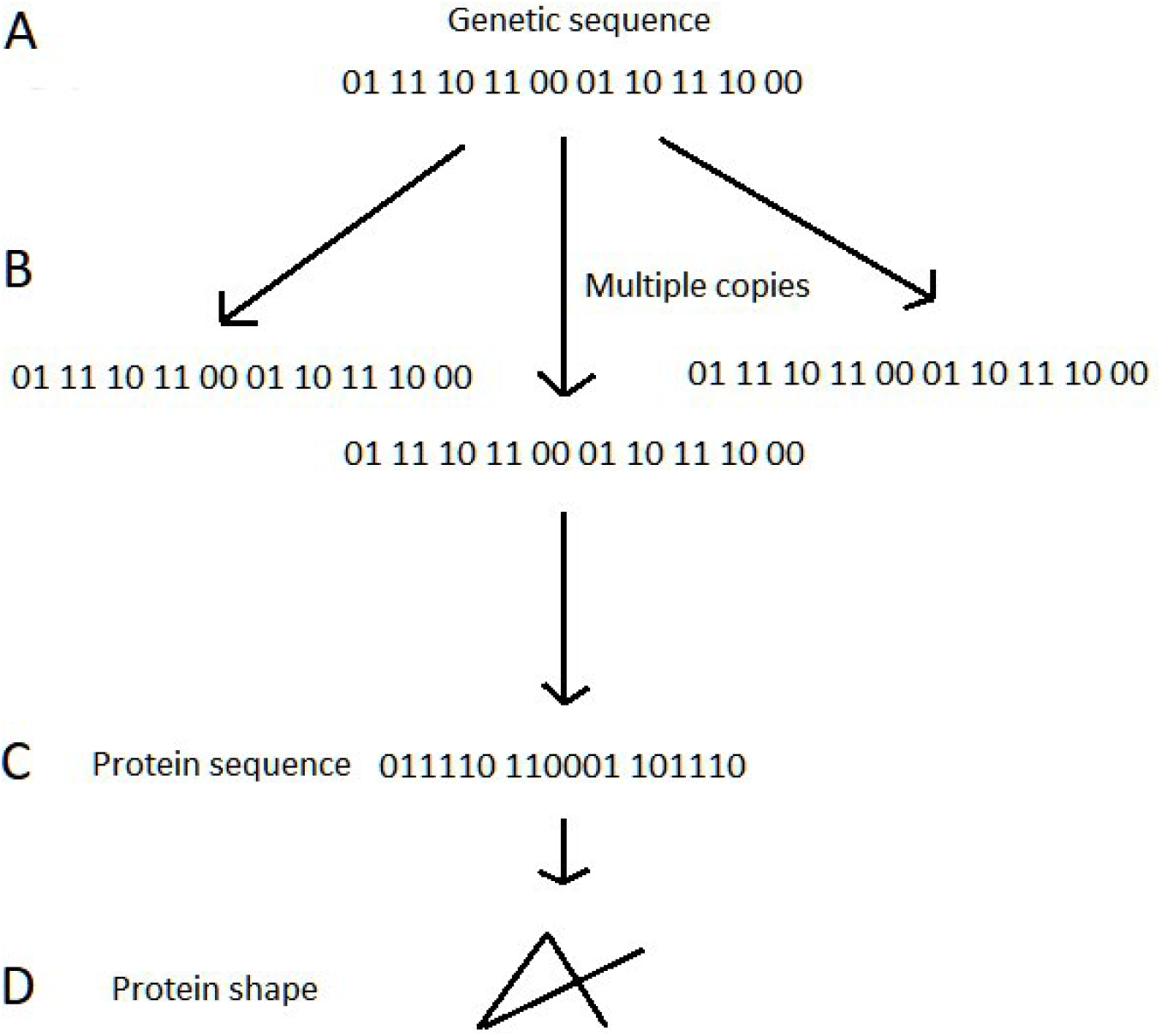
Simplification of the pathway for conversion of one dimensional genetic sequence to a three dimensional protein shape. A. Genetic sequence where each word is 2 bits in width. This sequence is representative of the proximate code for a protein molecule. B. Multiple copies of the genetic sequence are created from (A). C. Protein sequence is formed from (B), where each word of the genetic sequence is 6 bits in width, although the bit size is compressed in the protein code. D. Protein shape in three dimensions, where the shape is formed by the dynamics of physical interactions within the protein and between other molecules.

It is presumed that the processing of a linear sequence of genetic information to a protein shape is an optimized process, although constrained by the mechanisms of biological pathways. The linear sequence is one dimensional, a compression of the three dimensional shape of the protein, and its corresponding spatiotemporal dynamics in a biological system. Biological molecules are the operators to convert the information along the steps of this pathways, as an algorithm.

#### 4.2.3. Cellular immunity in jawed vertebrates

The immune system of jawed vertebrates has a similar strategy. In this case, the genetic information is not inherited, but originates from a foreign source, such as a virus or bacterium, and the infection may cause changes in the host. To deter foreign invaders, a jawed vertebrate host has cellular processes that search for intracellular and extracellular proteins, whether the source is foreign or from itself, for detection of pathogens by their proteins and to eliminate the infected host cells.

A virus is defined as a pathogen that is dependent on a host cell for genetic replication and the generation of its proteins (Fig. 4). After infection, the viral proteins are produced in a host cell, so these proteins and their sequence are available for detection by the host system. This involves a pathway which is similar to the information processing in the genetic inheritance of cellular organisms. The protein information, the sequence of amino acids, is processed to a set of subsequences. These subsequences are small in size as compared to the original sequence, and these are further combined with a specialized cell surface receptor [51–52]. This combination leads to a three dimensional shape, so the one dimensional protein subsequence has been converted to a three dimensional structure, and this latter arrangement is used in detection by specialized cells that surveil the surface of host cells [53].

**Figure 4.**
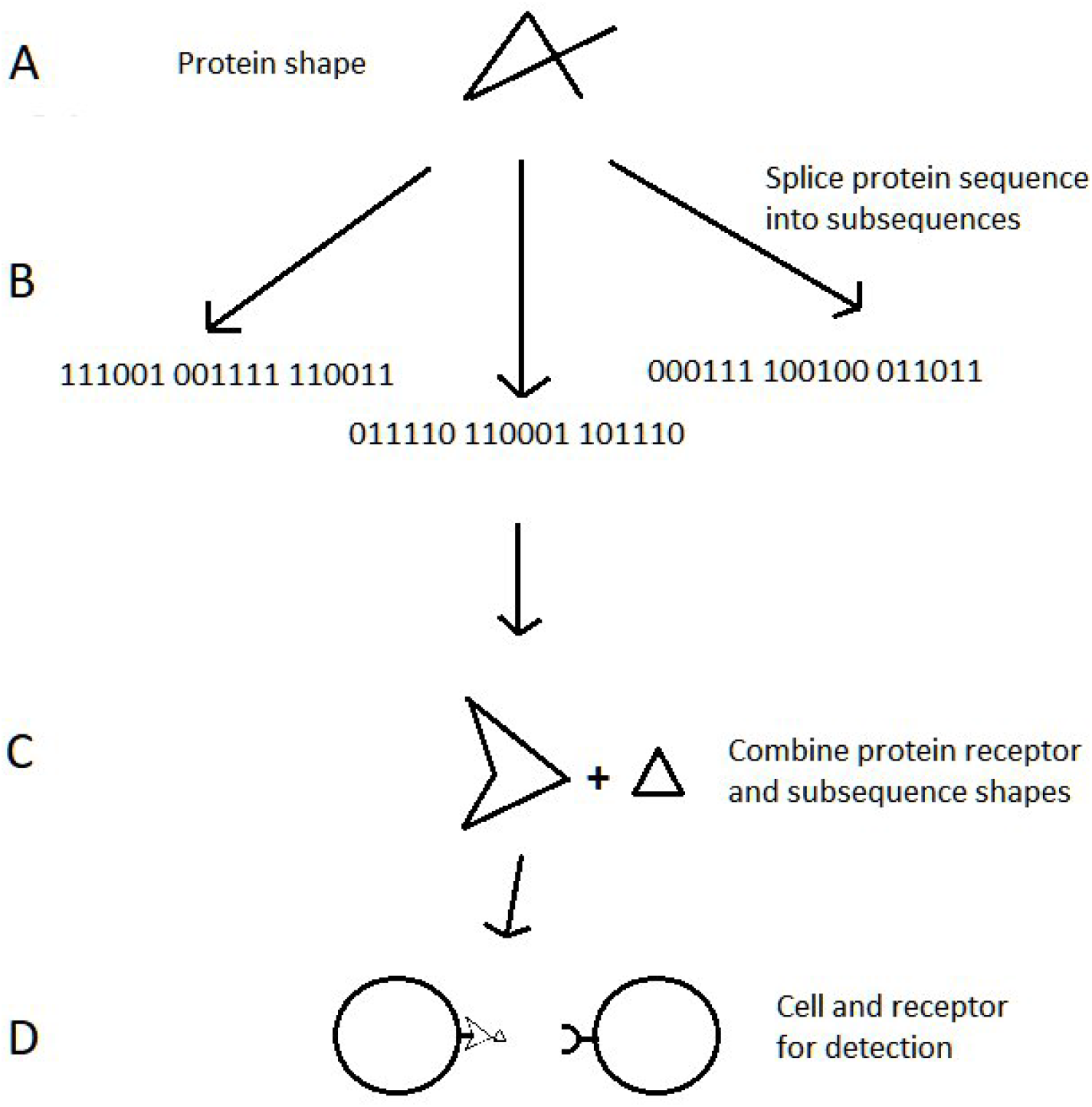
One simplified pathway of jawed vertebrate immunity, from intracellular pathogenic protein to its detection by a host cell. A. Shape of viral protein in three dimensions. B. Protein is spliced into protein subsequences. C. Subsequence of viral protein is combined with a host cell receptor. D. The protein subsequence and receptor from (C) are bound to the surface of a host cell (on the left). A specialized cell (on the right) with a cell surface receptor scans host cells for evidence of protein subsequences from a foreign source. Positive detection of a viral protein may contribute to an immune response, so that host cells infected with a pathogen are eliminated.

The above mechanism of cellular surveillance involves another type of cell surface protein receptor (Fig. 4D). This receptor is formed by an analogous informational process as described in the previous examples. It originates by different combinations of proteins that undergo a process of recombination and mutation, thereby modifying the protein sequence, and changing the resultant receptor shape, which leads to a very large diversity of receptor types as compared to the genetic instructions that code for the receptors [54]. This diversity of receptor types offers the host the possibility for detection of many kinds of pathogens, including those that are novel to the organism. This is immunity’s probablistic solution to the problem [55].

There is an additional layer of complexity in this process, where many different receptor types function in a complementary way in detecting pathogenic proteins. This overall process may be considered optimal in design, with high precision for detection of pathogens [54], while not reacting to proteins that originate from an uninfected host, because otherwise host cells without infection would be eliminated by the immune system [56].

The contest between survival of the host and the pathogen is dependent on the generation of the above protein subsequences by the host. The length of the relevant portion of these subsequences is typically short. For the intracellular case, they are often about 9 amino acids in length [54]. A short length of the detectable subsequence is expected. If the pathogen is continually and rapidly evolving by multiple mechanisms, such as mutation and recombination, then its genetic sequence is expected to eventually escape detection by the host immune system.

However, the evolutionary process is not likely to act on every portion of a genetic sequence, so subsequences are expected to stay unmodified over time. Furthermore, a type of pathogen, from a population perspective, is constrained in its evolvability, since genetic change may lead to loss of biological function, such as a change in the protein sequence which leads to a change in an essential portion of the three dimensional shape. This is a further constraint on the pathogen from evolving its genetic sequence beyond a host’s ability to detect it. Lastly, the protein subsequences are sufficiently long for detection precision by the host, otherwise, a very short subsequence is likely to originate from multiple sources, such as the host itself. This process is further regulated by its dependence on multiple and different subsequences for a strong immune response, and thereby the likelihood a pathogen is detected by an effective sampling process across the genetic sequence.

From the perspective of the pathogen population, its genetic changes may lead to an evasion of host immunity, such as in the case of the above intracellular pathway. It may avoid detection along the steps of this pathway (Fig. 4). One method is by interruption of the path as shown in Fig. 4B, so that a previously spliced protein subsequence is no longer spliced by the host. Another method is by interruption of the path in Fig. 4C, so that a spliced protein subsequence no longer robustly combines with the host cell receptor. A third method is by the path in Fig. 4D, so the specialized cell and its receptor no longer recognizes the combinational pattern on another host cell, where the pattern is presented as the combination of protein subsequence and cellular receptor. A population of a type of pathogen evolves evasion of host immunity by genetic changes, including recombination within and between populations.

From a higher level perspective, the host is in essence detecting the virus by sampling the viral protein subsequences, while the viral population is favoring genetic changes for evasion of host immunity. These kinds of favorable genetic changes are expected to change the corresponding protein sequence of the virus, and therefore increase the chance of evading host detection, but these particular changes are not expected to substantially alter the protein shape, otherwise there may be a loss of an essential biological function. This process appears optimal since the host is sampling genetic information of sufficiently high complexity for identifying pathogens, while avoiding detection of proteins of itself. In addition, the population of the host and pathogen may continue to interact over a long time scale since both have the potential to adapt to the other’s strategy, where the host population is learning the genetic sequence of the pathogen and the pathogen is evolving to evade this learning process. Persistence of this system depends on many factors, including population size and evolutionary rates, so it can be described as a high dimensional phenomenon.

#### 4.2.3. Neural systems in animals

Lastly, the neural network of animals can be considered as yet another example of an information-based system. It represents an algorithm that processes information as input and for output. For example, visual input is received across a two dimensional surface, but visual perception is in three dimensions. Therefore, the neural network is processing the information for generation of the perception [57]. Furthermore, the visual scene is processed into segments, for detection of objects and boundaries [58].

In cognition, the information processing is not by a biological enzyme, or by the strength of a protein interaction, but in the electrochemical-based communication of the neural network. To robustly recognize visual objects [59], this analog of an algorithmic system is expected to use the informational processes of combination, recombination, and other mechanisms, for generation of a large diversity of informational patterns for matching an observation to memory. As in the above examples, this system was formed by an evolutionary process that favors information optimality [60], while the neural system shares its network structure with the artificial neural networks from computer science, such as the deep learning systems [61–63].

The deep learning systems are capable of encoding the representations of information, allowing for a high dimensional processing, such as for representing the processes of the above biological systems. The gain in power to represent and predict a complex nonlinear dynamic system is offset by a loss in interpretability by use of the algorithm. A deep learning system can essentially serve as a mirror of Nature’s engine which drives information from an input state to that of output, where the engine mirrored is a path of processes that depend on the action of biological molecules [64–66].

